# Estimating the motor exploration in reinforcement learning

**DOI:** 10.1101/2025.01.21.634056

**Authors:** Anja T. Zai, Corinna Lorenz, Shakana Srikantharajah, Nicolas Giret, Richard H.R. Hahnloser

**Affiliations:** Institute of Neuroinformatics, University of Zurich and ETH Zurich, Switzerland; Neuroscience Center Zurich (ZNZ), University of Zurich and ETH Zurich, Zurich, Switzerland; Institut des Neurosciences Paris Saclay, UMR 9197 CNRS, Université Paris Saclay

## Abstract

What motor exploration strategies do animals use to learn a skill from rewards? Reinforcement learning theory provides no guidance for estimating motor exploration — the behavioral component aimed at discovering better strategies. Inspired by the brain’s modular organization, we postulate a latent learner that explores via an additive source of ideal randomness it injects into behavior.

Assuming the learner is ignorant of other motor components, which is sub-optimal by design, evolutionary fitness argues that these should display mainly non-ideal variability.

We test this recipe for behavior decomposition in songbirds subjected to a vocal pitch conditioning task. The explorative component we estimate from vocalizations accounts for the motor contribution of a basal ganglia pathway and the other behavioral component accounts for birds’ suboptimal learning trajectories. This congruence between normative exploration and brain organization suggests that the evolutionary pressure for behavioral optimality is lesser than to learn from purely random trials.

## Introduction

Reinforcement learning (RL) is a computational theory about identifying optimal behavior that maximizes the collected reward (Bhui et al., 2021; Sutton & Barto, 2018; Williams, 1992). RL is a successful strategy in games (Silver et al., 2018; Tesauro, Gerald, 1994) and holds promise as a theoretical framework for understanding neural processing, particularly in dopamine neurons (Hollerman & Schultz, 1998; Kim et al., 2020), but see (Jeong et al., 2022). Even though early concepts of RL were inspired by animal behavior (Sutton & Barto, 1981), applying RL to natural behaviors has remained challenging. Mainly, behavior tends to be sub-optimal, violating optimal action policies (Akaishi et al., 2014, p. 201; Akrami et al., 2018; Samuelson, William & Zeckhauser, Richard, 1988). Behavioral sub-optimality by itself does not falsify RL theory; it is the overestimated amount of exploration that remains perplexing.

Exploration is a central component of RL; it is conceived as actions that deliberately deviate from an optimal strategy in the hope of finding a better strategy leading to more reward, also known as the exploration-exploitation tradeoff. RL theory applies to the entire range from pure exploitation to pure exploration (Watkins & Dayan, 1992) and so there are virtually limitless ways of introducing exploration into RL models, i.e., the theory is under-constrained and offers no principled way of estimating the behavioral component attributable to exploration. Despite this modeling freedom, when fitting behavior with normative RL models that incorporate an exploitation-exploration tradeoff, subjects seem to learn less from deviant behavioral variants than they theoretically could (Drugowitsch et al., 2016; Niv, 2009), which means that some of their behavioral variability stems not from decisions to explore (Findling et al., 2019).

Indeed, there is growing evidence that the brain is capable of learning from only a part of motor variability (Dhawale et al., 2017; Therrien et al., 2016) and that some sources of motor variability are disconnected from a learning experience (Therrien et al., 2016). For example, motor noise is thought to provide little benefit for improving behavioral output (van Beers, 2004), and artificially imposed motor noise is of no benefit to subjects for increasing the success of their actions (Chen et al., 2017; Therrien et al., 2018). A behavior-congruent RL theory therefore needs to account for more than just exploitation of good strategies and exploration of new strategies.

## Results

To keep the mathematical rigor of RL but to make it amenable to natural behaviors and their hidden explorative components, we abandon the overall behavioral optimality as a goal of RL. Instead, we frame RL as a model of a hypothesized latent learner, a behavioral module that learns from its random perturbations of behavioral output. Such a behavioral module dedicated to explorative motor variability has been hypothesized (Wirthlin et al., 2019) and it encapsulates that many skills are learned and recalled in separate brain regions (Andalman & Fee, 2009; Kawai et al., 2015; Kojima et al., 2018; Olveczky et al., 2005). To formulate latent RL as a normative principle suitable for statistical inference (Młynarski et al., 2021), we assume that the latent learner maximally learns from the effects of its perturbations on reward, whereas all other modules are unable to meaningfully integrate reward experience on a trial-by-trial basis.

We assume that the latent perturbations exclusively form an ideal source of randomness, i.e., the latent learner alone (exclusively) contributes to behavioral trials in terms of independent and identically distributed (iid, ideal) random perturbations. We motivate this assumption of ideal exploration with computational generality and with evolutionary fitness: On the one hand, random sampling is the basis of science as it guarantees the collection of unbiased data. For example, randomness in clinical trials avoids spurious correlations with uncontrolled influences. In games such as rock, scissors, and paper, random strategies make a player robust to adversarial counter strategies. On the other hand, the ‘exclusivity’ attribute, i.e., that no other module contributes iid additive motor variability, maximizes the agent’s learning efficiency (think about the converse in the context of randomized trials: when you ignore whether you administered a treatment or a placebo to a subset of subjects, your effect size will decrease).

Latent RL offers a simple principle for dividing behavior into three components (Fig. 1a):

**Figure 1:**
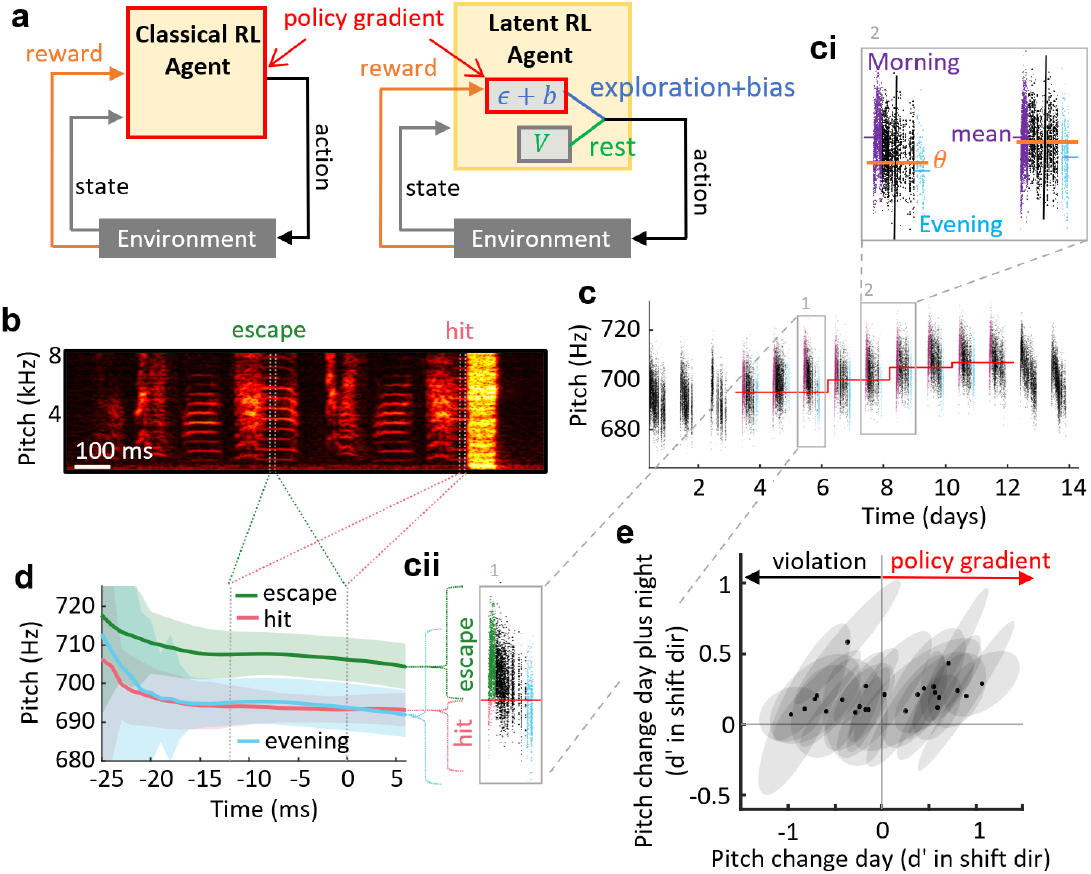
Latent RL is designed to overcome behavioral incongruences of classical RL. **a** In classical RL (left), an agent potentially modifies all behaviors to increase reward — the agent senses the entire policy gradient. In latent RL (right), policy gradient search is restricted to a latent learner that generates ideal explorations ϵ to learn a reward-increasing motor bias b. The learner ignores the agent’s non-ideal behavioral primitives V(rest). **b** Song spectrogram (top) of a bird subjected to aversive WN stimuli (hit, yellow area) contingent on low-pitch syllables. The pitch measurement window is indicated by vertical white dotted lines. **c** Pitches (dots) of the targeted syllable before, during, and after contingent WN exposure (the red line delimits the pitch threshold θ for WN playback). **ci** Evening pitches (blue dots) are lower than morning pitches (purple dots) despite the aversive stimuli they trigger. **cii** Evening pitches (blue) are closer to WN-eliciting pitches in the morning (hits, red), than to WN-escaping pitches. **d** Average pitch trace as a function of time in the syllable (± 1 std in shading). The bird uniformly (in time) changes pitch towards the previously penalized (hit) syllable. **e** In half of birds, the morning-to-evening pitch change (x-axis) is opposite to the morning-to-morning pitch change (y axis, dots represent single birds and shaded ellipses represent 1 std): All birds respond aversively to the WN stimulus (y>0), but half of them learn non-monotonically (x<0), in violation of an agent-level policy-gradient strategy.

1. Explorative ideal perturbations *ϵ*,
2. their optimal reinforcement (a learned motor bias *b*, also part of the latent learner), and
3. all remaining behavioral patterns *V* the agent can generate (the rest).

To work out, a latent RL model must produce good fits to behavior and it must do so self-consistently, i.e., the perturbations estimated from data must be close to ideal. This latter property can be tested via mutual information, a quantity that is minimized by independent (ideal) random variables.

We apply this idea to songbirds while we instrumentally condition their song away from the stable version they acquired from a tutor (Tumer & Brainard, 2007), a faculty that depends on a cortico-basal ganglia (CBG) loop (Ali et al., 2013; Andalman & Fee, 2009; Charlesworth et al., 2011, 2012; Hoffmann et al., 2016; Zai et al., 2020) and its output, the lateral magnocellular nucleus of the anterior nidopallium (LMAN). As conditioning stimulus, we play a loud white-noise (WN) stimulus through a loudspeaker when the pitch (fundamental frequency) of a chosen song syllable is below (or above) a set threshold, Fig. 1b (see Methods).

Zebra finches that we subjected to this paradigm escaped the aversive reinforcer (penalty) by adapting the pitch of the targeted syllable away from the hit zone (Fig. 1c), against their tendency to maintain a stable pitch. Birds coped with the tradeoff to maintain a stable baseline song and to escape the penalty by violating traditional notions of optimality in RL: In *n* = 10/18 birds, the pitch dynamics were non-monotonic; these birds did not escape WN during the day, but they did so over the course of 24-h periods, Fig. 1c-e. Such non-monotonicity clashes with earlier conclusions that learning results from reinforcement of successful past actions (Charlesworth et al., 2011), since these birds seemingly just did the opposite, which is to selectively repeat the penalized syllable variants. Non-monotonic learning argues against pure (policy-gradient) RL models of reward maximization (or punishment minimization) and so it argues against classical RL theory.

To align RL models with these diverse empirical pitch adaptation strategies, we designed a latent learner (Fig. 2a) that:

**Figure 2:**
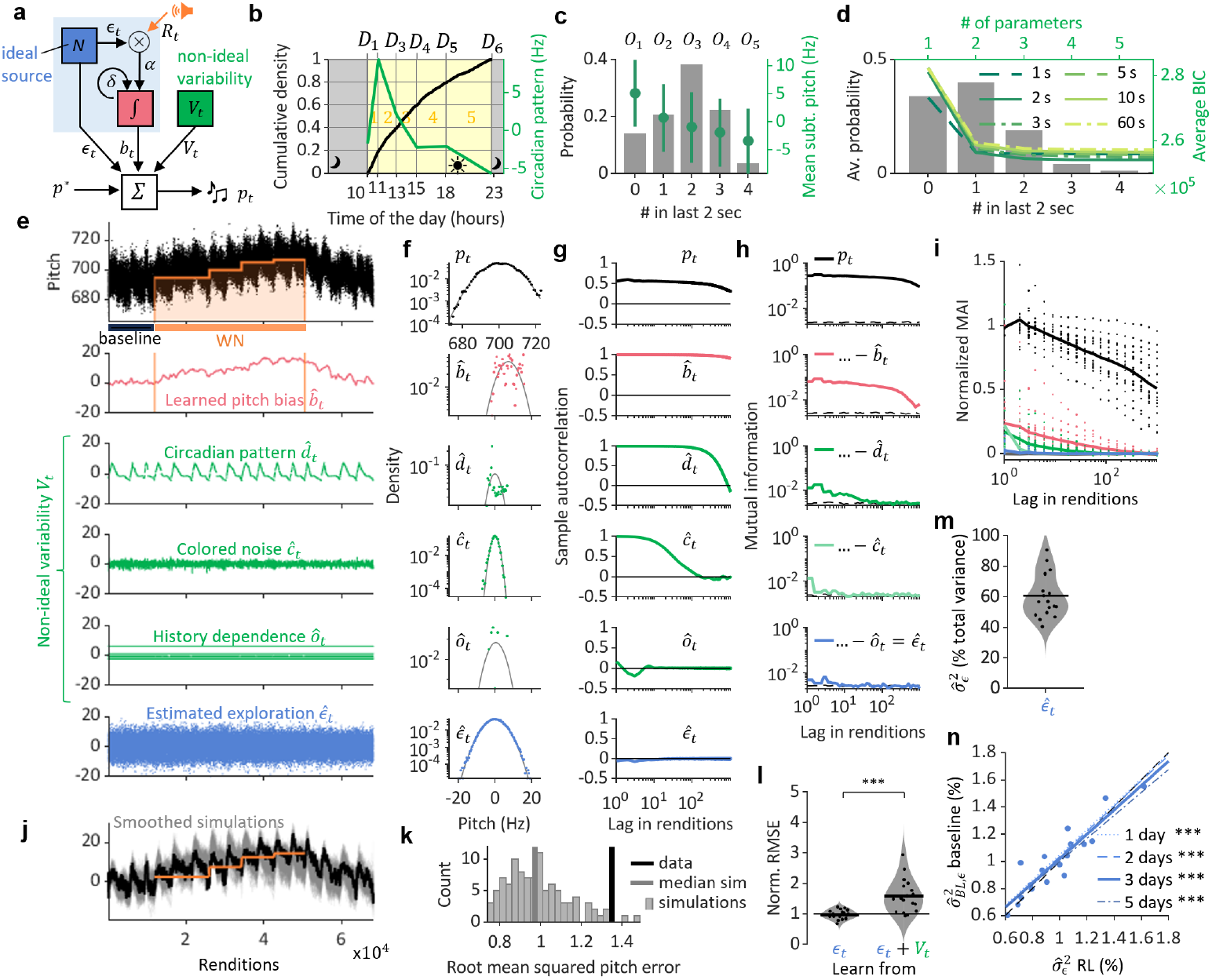
Latent RL self-consistently fits diverse learning trajectories. **a** We simulate a latent learner that produces explorations ϵ_t_ to increase reward R_t_ via a motor bias b_t_ that shifts the pitch p_t_ away from its target p^*^. The agent also exhibits non-ideal behavioral components V_t_ (green) composed of a circadian pattern d_t_, slow drift c_t_, and history dependence o_t_. **b** The circadian pattern (green) we modeled as a piecewise linear function defined by six coefficients D_1,…6_ spread across 5 daytime bins (yellow shading). Each bin contains 20% of syllable renditions (illustrated by the black normalized cumulative distribution; same bird as in Fig. 1, baseline period). **c** The history dependence we modelled as mean-subtracted pitch offsets O_k_ (±std, green), where k = 1, …, 5 is the number of syllable renditions within the last Δ = 2 seconds (gray bars). **d** The average Bayesian information criterion (green lines, N = 18 birds) was lowest for Δ = 2 seconds (green solid line). **e** After model fitting, the pitches p_t_ (black dots, top) are additively decomposed into the components: 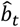 (bias, red), V_t_ (green), and the estimated exploration 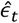(blue) with distributions shown in **f** and sample autocorrelations in **g**. Same bird as in d. **h** The mutual average information (MAI) of the pitch residuals if formed by cumulatively subtracting from the pitches p_t_ the estimated behavioral components (top to bottom). The MAI of the ultimate residual (blue, bottom) corresponding to the estimated explorations 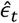 is commensurate with sampled white noise (dashed). **i** Same is h but shown as averages (lines) across N = 18 birds (dots represent individual birds). **j** The observed pitch trajectory lies within 100 simulated trajectories (gray). **k** The histogram of root mean-square pitch errors (RMSEs, gray) between simulated trajectories and their median. The RMSE of the observed trajectory (black) is larger than the median error (gray line), but within the stimulated distribution of errors. b-h, j, k: same bird as in Fig. 1. **l** The RMSEs normalized to the median simulations in N = 18 birds (dots). The average RMSE of latent RL is consistent with data, it equals E_n_ = 1.0 (horizontal bar, left), unlike classical RL with an average RMSE of E_n_ = 1.8 (right bar, *** p < 0.001, paired t-test). **m** The relative magnitude of exploration calculated as the estimated variance of exploration 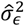 divided by the baseline pitch variance (N = 18 birds, dots). **n** The estimated standard deviation of exploration 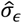 in percent of p^*^ generated by the latent learner (blue dots, N = 18 birds) during RL (x-axis) and during baseline (y-axis). Linear fits (blue lines, *** represents p < 0.001) for different numbers of baseline days used (legend), the fits are close to the diagonal (black dashed line).

1. generates Gaussian iid trial-by-trial pitch perturbations (ideally perturbs),
2. correlates the perturbations with the presence/absence of WN (correlates), and
3. leakily integrates the perturbations into a motor bias (integrates).

The correlations (2) detect rewarded (unpunished) perturbations, their integration (3) constitutes the reinforced pitch bias, and the leaky integration (3) models the attractive pitch target (the stable learned song) under absence of WN. There is broad experimental support for computations 1-3 in the songbird AFP with its dopaminergic recipient Area X and its output nucleus LMAN:

- In support of ideal perturbations (1), suppressing LMAN and Area X activity in both juveniles and adults greatly reduces the trial-by-trial variation of song (Kao & Brainard, 2006; Olveczky et al., 2005; Singh Alvarado et al., 2021).
- In support of correlation (2), dopaminergic input feeding into the AFP is required for pitch adaptation (Hoffmann et al., 2016), and the contingency between dopamine release and motor output sets the direction of the learned pitch bias (Hisey et al., 2018; Xiao et al., 2018).
- In support of integration (3), after lesions in the AFP, adult birds lose the ability to adapt pitch (Ali et al., 2013); and inactivation studies reveal that LMAN biases song pitch to avoid penalties (Andalman & Fee, 2009; Warren et al., 2011).

This latent learner is characterized by mainly 3 parameters that we seek to infer from observed pitch trajectories: the (exploration) variance 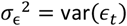 of its iid perturbations *ϵ*_*t*_ (at time *t*), the reinforcement learning rate *α*, and the leakiness *δ* of the bias integrator (see Methods). This simple model encapsulates an optimal agent that however by itself produces poor fits to the behavioral data and is not self-consistent: Although the inferred explorations often were roughly Gaussian distributed, they were far from ideal, as evidenced by their large mutual average information (MAI) exceeding that of ideal Gaussian white noise by about one magnitude on average (see Methods, Supplementary Fig. S1).

After some search of behavioral components that make the latent learner self-consistent, we obtained good results using three behavioral components (see Methods for mathematical definitions, Fig. 2):

1. a circadian pitch pattern *c*_*t*_,
2. a slow random drift *s*_*t*_, and
3. a pattern of history dependence *o*_*t*_.

The circadian pattern (Fig. 1) we modeled as a piecewise linear function (with time bins each containing 20% of syllable renditions, Fig. 2b), the slow drift as non-iid colored noise (i.e., not part of the learner module), and the pitch history dependence we modeled in proportion to the number of target syllables sung within the last 2 seconds (Fig. 2c,d). The estimated model components (Fig. 2e) each contributed considerably to total pitch variance (Fig. 2f) and sample auto-covariance (Fig. 2g). Model self-consistency was evidenced by the estimated explorations being Gaussian distributed (Fig. 2f) and having roughly iid statistics, as shown by the average lag-dependent MAI (Fig. 2h) that on average was only 13% above that of pure white noise (Fig. 2i).

The self-consistent models produced good fits to observed pitch trajectories and they generated trajectories that were like the observed ones in terms of the root mean-square pitch error (RMSE) that we used to compare observed with simulated trajectories (see Methods, Fig. 2k, l): The normalized RMSE *E*_*n*_ (the observed RMSE in each bird divided by the mean RMSE of simulations) averaged to *E*_*n*_ = 1.0 across birds, implying that we could not reject the hypothesis that the observed pitch trajectories were generated by the model (*p* = .7, t-test for *H*0: *E*_*n*_ = 1, *df* = 17), unlike classical RL models that learned from the entire behavioral variability (*E*_*n*_ = 1.8, *p* = 2 * 10^−4^, t-test for *H*0: *E*_*n*_ = 1, *df* = 17, see Methods). Only in one of 18 birds was the observed RMSE larger than 99% of simulated RMSEs, Supplementary Fig. S2. These results reveal that overall, unlike in classical RL, the errors made by latent RL models fits were consistent with the presumed ideal variability.

The estimated exploration variance 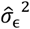(indicated by 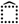) during WN we expressed as a fraction 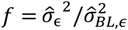 of the total baseline (BL) variance 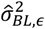 was quite large, averaging to 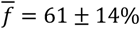 (*n* =18 birds, Fig. 2m). The broad range of *d* from 41 to 91% (*n* = 18 birds) suggests that animals differed in their propensity to explore over a factor of more than two (or, put differently, they differed by a factor of two in their magnitude of non-explorative behavioral components).

We estimated explorations also during the preceding baseline period, when the pitch dynamics were similarly complex as during the subsequent conditioning period. The associated baseline exploration variance 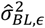 was highly correlated with the WN exploration variance 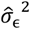 (Pearson correlation coefficient *ρ* = 0.91, *p* = 1.4 * 10^−7^, *N* = 18 birds, see Methods, Fig. 2n). The slope *m* of a linear fit of 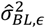 as a function of 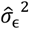 was not different from the unit slope (*m* = 1.02, *p* = 0.5, *N* = 18 birds, t-test), suggesting that zebra finches do not modulate their exploration variance during song practice, even while subjected to WN penalties.

Explorations and their variance could be estimated from few renditions only. The estimated baseline exploration variance 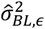 from half a day of song data was correlated with 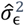 at *ρ* = 0.87 (*p* = 3.1 * 10^−6^, *N* = 18 birds) and even when estimated from merely 100 syllable renditions, the baseline exploration variance was correlated at *ρ* = 0.80 (*p* = 6.7 * 10^−5^, *N* = 18 birds). That it is possible to estimate explorations from so few examples makes the model very versatile.

We further tested latent RL as a plausible neural mechanism, i.e., whether the learner module captures the role of a motor-learning pathway. Pitch adaptation in songbirds depends on a cortico-basal ganglia loop: Without its output nucleus (LMAN), song variability is largely reduced (Kao and Brainard, 2005) and adult birds fail to learn from aversive pitch reinforcement (Charlesworth et al., 2012). We thus hypothesized that the estimated explorations match with the pitch variance contributed by LMAN.

We modeled bilateral lesions in LMAN (Fig. 3a) by setting both pitch explorations and the learned pitch bias to zero (Fig. 3b). Lesions did not affect the overall spectral and temporal structure of song motifs (example in Fig. 3c), but they decreased the average daily pitch variance by 22 ± 25% on average (Fig. 3c,d,f), in line with previous reports (Hampton et al., 2009; Kao & Brainard, 2006; Thompson et al., 2011). Across birds, lesion extent was moderately correlated with the reduction in observed pitch variance (Pearson correlation *ρ* = 0.54, *p* = 0.01, *N* = 20 birds, regression fit explains 33% of variance, Fig. 3f). In contrast, lesions reduced the estimated exploration variance 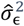 by 40 ± 32% on average and lesion extent was strongly correlated with exploration decrease (ρ = 0.84, *p* = 2.1 * 10^−5^, regression fit explained 70% of the variance Fig. 3f). Thus, exploration was a better predictor of LMAN lesion outcome than was pitch variability. We found that exploration was also a better predictor of lesion outcome than a simpler model (Toutounji et al., 2024) in which LMAN’s effect is estimated as the motor residual following pitch detrending (*R*^2^ = 0.45, *p* = 0.001, *N* = 20 birds). Thus, neither pitch nor detrended pitch capture LMAN function as well as does pitch exploration, in support of our hypothesis that LMAN generates ideal pitch explorations.

**Figure 3:**
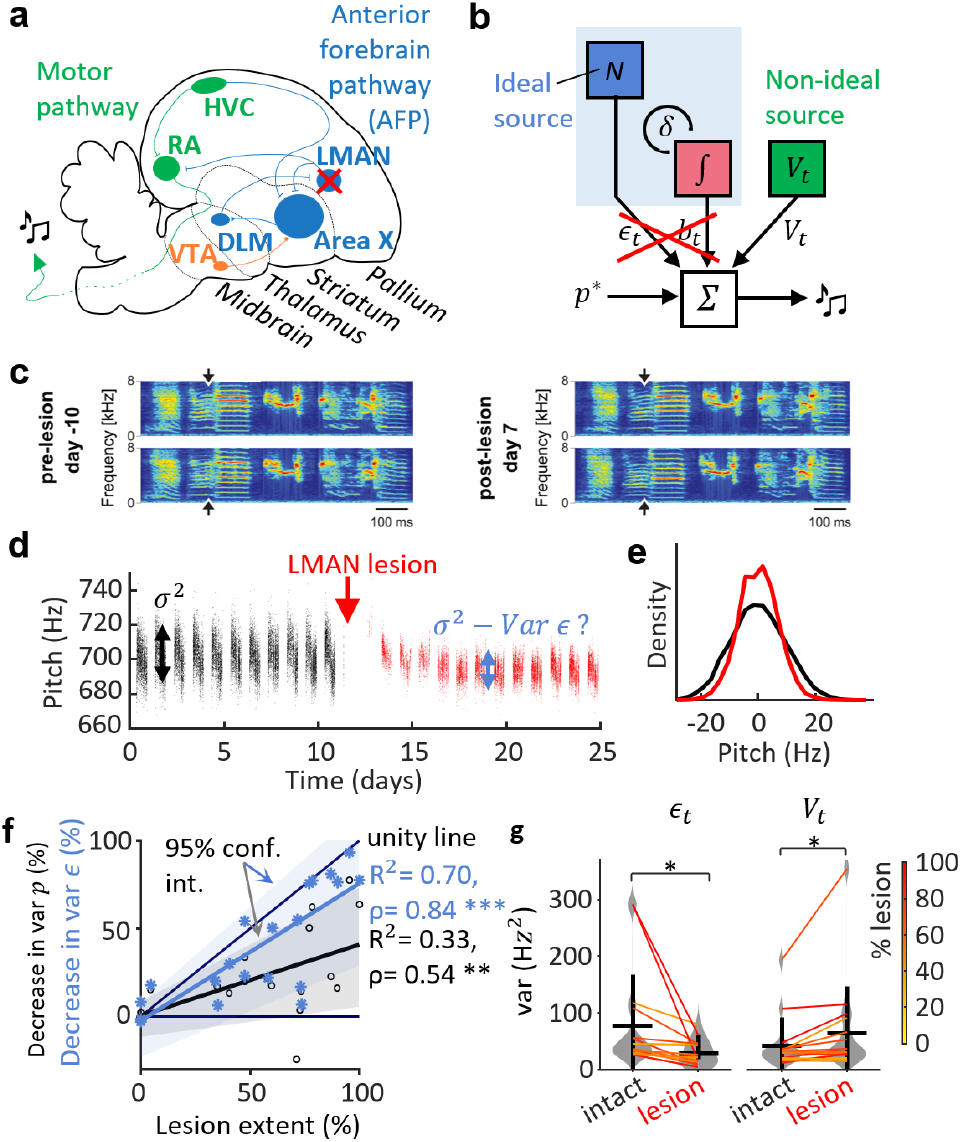
LMAN lesions eliminate explorations. **a** Brain schematic of the song system. We bilaterally lesioned the lateral magnocellular nucleus of the anterior nidopallium (LMAN). **b** We hypothesized that LMAN lesions abolish both explorations ϵ_t_ and the learned pitch bias b_t_. **c** Two example motifs before (left) and after (right) LMAN lesions. The lesions did not degrade the spectro-temporal motif structure. The arrows indicate the window measurement window of the target syllable. **d** Pitch trajectory of an example bird showing reduced pitch variability (red) following LMAN lesion; **e** Normalized histogram (density) of mean-subtracted pitch before (black) and after (red) LMAN lesion (same bird as in d). **f** The estimated exploration variance correlates better (blue, N = 20 birds) with lesion extent than does the decrease in pitch variance (black). Shown is the type-1 linear regression and 95% confidence interval estimated from 100 bootstrapped replicates. The reported statistics are the Pearson correlation coefficient ρ and the percent explained variance R^2^, * p < 0.05, and *** p < 0.001. **g** Change in exploration variance before and after LMAN lesion (left) and change in variance of remaining behavioral components V_t_ (right, N = 17 birds with non-zero lesion volume).

We found that our approach also holds promise in human subjects that were instructed to repeat a simple utterance while trying to avoid a WN stimulus that we delivered contingently on their pitch (see Methods). During WN sessions, subjects successfully changed the pitch away from the WN zone and after cessation of WN delivery they tended to revert their pitch back towards baseline (Fig. 4, see Methods). A simple latent RL model that included merely an additional colored noise source proved sufficient to self-consistently estimate subjects’ pitch explorations, Fig. 3h. The estimated exploration variance 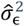 during WN delivery was correlated with the exploration variance during baseline (correlation coefficient *ρ* = 0.63, *p* = 0.017, *N* = 14 human subjects). However, the slope *m* = 0.78 of a linear fit of the WN vs baseline variance was different from unity (*m* = 1.0, *p* = 0.01, t-test, *N* = 14), suggesting that subjects increased the exploration variance during WN delivery (active inference).

**Figure 4:**
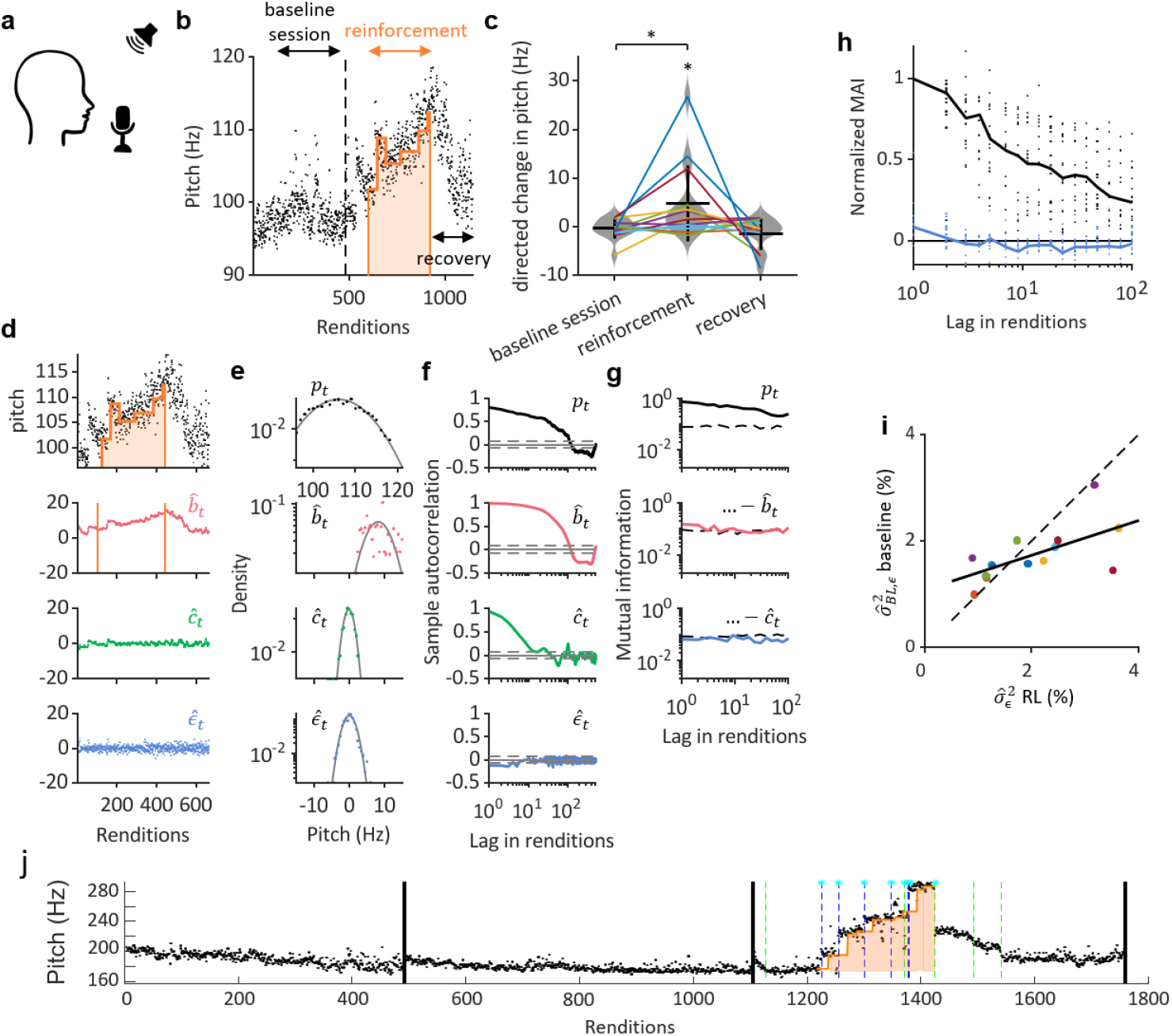
Pitch conditioning in human subjects. **a** Subjects were instructed to repeat the utterance ‘da’ at a rate of roughly twice per second. Same pitch paradigm as in birds. **b** Pitches (dots) of ‘da’ before, during, and after conditioning, in an example subject. During the baseline session, no white noise is played. During minutes 2-5 of the WN session (delimited by the orange curve), aversive reinforcement (a loud white noise) is played upon low pitch renditions. During recovery, no white noise is played. **c** Pitch change (away from the white-noise zone, e.g. up in b) during the baseline session, during the reinforcement period, and during recovery (N = 14 subjects indicated by lines, * p < 0.05, t-test). **d** After model fitting, the pitches p_t_ (black, top, same subject as in b) are additively decomposed into a bias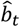 (red), a colored noise 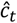 (green), and the estimated exploration 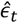 (blue) with distributions shown in **e** and sample autocorrelation in **f. g** The mutual average information (MAI) of the residuals (top to bottom, formed by cumulatively subtracting the indicated behavioral components) from the pitches p_t_ are gradually reduced. The MAI of the estimated perturbations 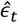 (blue, bottom) is as low as that of sampled white noise (dashed line). **h** The normalized MAI of pitch trajectories (black dots, N = 14 subjects) and of estimated perturbations (blue dots, N = 14 subjects), and their averages (lines). **i** The estimated explorations 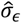 in percent of p^*^ (colored dots, N = 14 subjects) are significantly correlated between baseline (no RL, y-axis) and RL (x-axis) periods (linear fit: solid line, p = 0.001, Spearman’s correlation; identity: dashed line). **j** Subject who increases the pitch step rate in the direction of WN escapes (6 of 8 steps during WN are aligned with the WN escape direction, Z = 2.0). Pitch steps preceded by WN are indicated by a cyan asterisk, up steps by dashed blue lines, down steps by dashed green lines).

Humans differ from zebra finches in that the former can dramatically change vocal pitch from one rendition of an utterance to the next (Zai et al., 2024), a possible expression of stepwise exploration or insight learning (a form of exploitation). Indeed, in zebra finches we found no pitch discontinuities or steps where past and future pitch averages would differ by more than Θ = 5% (see Methods). However, in 14 human subjects we found a total of 103 pitch steps (Θ = 5%, mean: 3.6 ± 4.6 (std) steps per subject, range 0 − 14 steps). When eliminating the discontinuities (see Methods), the correlation between WN and baseline exploration variance remained high (*ρ* = 0.71, *p* = 0.005, *m* = 0.77). Correlated WN and baseline exploration variances were robustly observed for a wide range of thresholds (Θ = 1%, 10%), showing that ideal exploration estimation is robust to pitch steps.

Pitch steps could constitute a human exploratory strategy on their own, since among the 103 pitch steps, 51 (50%) were preceded by recent WN experience (in the last 20 utterances), many more than the chance level of 20% defined as the fraction of utterances with preceding WN (test for equal proportions, *p* < 10^−24^, see Methods). Five of fourteen subjects significantly changed their pitch stepping behavior in response t WN. All five subjects produced significantly more pitch steps during WN than during baseline (test of equal proportions, *p* < 0.05, see Methods), suggesting that they adopted a pitch stepping strategy to explore the contingency of WN penalties, reminiscent to Levy flight foraging (Viswanathan et al., 2008).

Individual subjects produced too few pitch steps during WN (mean 3.6, range 0-14 steps) to test whether they learned also to produce pitch steps as an exploitative strategy, with exception of one subject who seemingly used pitch steps to escape WN (6/8 steps during WN were directed towards escape, *Z* = 2.0, Fig. 4j). Thus, although not all subjects seemed to make use of pitch stepping, some seemed to use it as a directed vocal exploration strategy and one managed to skillfully use stepping to escape WN. Pitch stepping harbors the obvious benefit of serving to explore more distant rewards in fewer trials (steps). Also, by being flexibly engageable (i.e. during WN), stepping is no obstacle to skilled stereotyped behavior.

## Discussion

To overcome the misalignment between RL theory and animals’ action choices, we have introduced a latent learner that agrees with a wealth of published data and that produces excellent fits to behavior even during sub-optimal (non-gradient-like) learning periods. By defining exploration as ideal motor perturbations, its impact on behavior can be estimated on a trial-by-trial basis, both in task and task-free settings.

We abandon unrealistic notions that behavior must be optimal and fully compliant with knowledge seeking. By restricting exploration to a learner’s ideal perturbations, we allocate optimal (gradient-following) learning to a distinct module rather than to the learning agent itself. This inherent optimality makes latent RL a normative theory, a framework for disentangling the components of a motor gesture, in particular the component that interacts on a trial-by-trial bases with external reinforcement.

The iid statistics of exploration are assumed to hold in the sensory pitch space rather than in a peripheral (muscle) space where independent Gaussian noise tends to be inefficient (Chiappa et al., 2024). We recognize that other sources of behavioral noise may be approximatively iid as well, but the presumed evolutionary efficiency led us to assume that their magnitude be small (i.e., neglectable), especially during skilled behavior where variability is inversely related to fitness.

Exploration predicts basal ganglia function better than does behavioral variability. Previous basal ganglia models have implicitly assumed that all of motor variability is exploration (Brudner et al., 2023; Toutounji et al., 2024), including models that refer to exploration as noise (Sankar et al., 2022). The idea of distinguishing exploration from noise is not new (Dhawale et al., 2017; Therrien et al., 2016), our work merely contributes a general means for estimating exploration.

By sub-optimal design, the latent learner deliberately dismisses some information such as motor efference copies (that are dispensable for generating pure randomness). This assumption clashes with suggested existence of an efference copy in LMAN of pitch variability generated elsewhere (Charlesworth et al., 2012), but it agrees with a more recent proposal that no efference copy in LMAN might be needed, provided that LMAN is informed about an auditory target that it can translate into a directional motor command (Zai et al., 2024). In that vein, we imagine that the learner’s output, for example its bias, is influenceable by upstream modules akin to directed exploration.

Latent RL differs from previous work in that it assumes that ideal exploration is not merely sufficient, it is necessary. For sufficiency of iid exploration there is broad support, since iid (e.g. Gaussian white) motor variability is a standard model component (Albert et al., 2021; Churchland et al., 2006; Dhawale et al., 2017, 2019; Therrien et al., 2016, 2018; van Beers, 2007). The iid assumption is usually motivated by convenience rather than as a criterion of self-consistency. Likely, assumed iid variability produces good fits because human motor adaptation is usually studied during short periods with few trials (Izawa & Shadmehr, 2011; Shmuelof et al., 2012), a setting like ours (Fig. 4) in which sub-optimal learning trajectories might not necessarily show up. That ideal exploration is also necessary is less obvious but is suggested by a recent study on the microstructure of learning in humans. (Findling et al., 2019) attributed non-optimality of human behavior to a noise source that engendered behavioral correlations across successive decisions, i.e., a non-ideal source in our terms. Whereas these authors attributed behavioral non-optimality to computational noise, we attribute it more broadly to arbitrary behavioral modules such as circadian fluctuations that contribute non-ideal variability and that are uninformed about trial-by-trial reward.

There are diverse mechanisms the brain could use to produce random explorations and maintain their independence. Randomness generators have been proposed on multiple spatial scales ranging from individual ion channels, to spontaneous synaptic vesicle release, to recurrent neural networks under inhibitory-excitatory balance (Darshan et al., 2017; Destexhe et al., 2003; Van Vreeswijk & Sompolinsky, 1996). On top of that, ideal perturbations could be promoted by synaptic learning rules that act on mutual information (Isomura & Toyoizumi, 2016; Toyoizumi et al., 2004), in line with the brain’s strategy to remove temporal correlations (Salisbury & Palmer, 2016).

Ideal statistics of exploration perfectly aligns with the brain’s sensitivity to sensory errors and its ability to modulate error sensitivity (Albert et al., 2021; Herzfeld et al., 2014). Namely, self-generated perturbations of behavioral output cause fluctuations of sensory feedback (sensory errors). But when perturbations are ideal, consecutive fluctuations are uncorrelated in sign and so do not affect sensitivity to sensory errors (Albert et al., 2021). Thus, ideal perturbations by themselves do not trigger known mechanisms for adapting to external perturbations. The sensory apparatus thus seems to filter out self-generated motor fluctuations with ideal statistics, presumably to increase sensitivity to correlated errors that are externally driven. Ideal experimenting therefore promotes adaptability in a dynamic world, perhaps in the service of robustly finding the true causes of increasing reward.

The zebra finches in our experiment seemingly did not modulate the magnitude of ideal exploration, but the additive nature of self-perturbations would make it simple to do that given a suitable timing signal (Fee, 2014). Past work suggests existence of exploration modulation. Motor variability can depend on feedback (Izawa & Shadmehr, 2011) and it can be regulated by both fast and slow reward-dependent processes (Dhawale et al., 2019). Zebra finches can recruit variability to direct learning to a subset of motor gestures (Ravbar et al., 2012) and they change variability with behavioral context, e.g. singing more stereotyped songs directed to a female, which is accompanied with reduced bursting in LMAN neurons (Kao et al., 2008). It remains to be seen which of these modulations of variability can be ascribed to exploration. Our reduced paradigm of singly housed birds seemingly did not elicit these modulations, but conceptually, latent RL can deal with such modulation as seen in humans, Fig. 4i.

On the one hand, large explorations allow for reaching more distant targets and rewards and so seem desirable — however note that large explorations do not necessarily imply faster learning, since learning speed in our model also depends on the learning rate unlike (Therrien et al., 2016, 2018). On the other hand, large perturbation may be a handicap, since the brain often learns more from small errors than from large ones (Marko et al., 2012; Robinson et al., 2003; Soetedjo et al., 2009; Wei & Körding, 2009). Humans seem to bypass the pitfalls of large ideal explorations by flexibly recruiting pitch stepping as an exploration mechanism, which needs further investigation.

In conclusion, latent RL holds promise as a simple strategy for building complex motor programs from simpler ones. Unlike classical RL, the challenge in latent RL is to transfer the knowledge gained from self-perturbations to motor primitives and more complex programs that control behavior in non-random ways. In our model, latent RL merely provides a simple bias towards more rewarding behavioral variants but the learner module is oblivious of how to meaningfully accumulate and consolidate the bias into extensive motor programs and motor plans. In this sense, latent RL can be seen as part of a more general perturbator-consolidator architecture (rather than an actor-critic architecture). In songbirds, the learned bias generated by LMAN is consolidated in HVC and the robust nucleus of the arcopallium (Tachibana et al., 2022). Processes of consolidation could be conceptualized in future work as a transfer of the learned bias to the repertoire of behavioral primitives, which offers large flexibility for incorporating ideal exploration into more elaborate models of behavioral learning.

We emphasize that latent RL does not preclude higher-level conscious learning mechanisms of choosing directed exploration strategies such as pitch stepping. In our view, latent RL is an evolutionary old adaptation that encapsulates the type of unconscious motor learning that is effortless and comes for free with practice. Latent RL can be used as a method for screening highly plastic subjects with large learning potential and may even be used for diagnosing learning deficits. Latent RL could be a promising route towards computational psychiatry, aiming to capture inter-individual behavioral variability in a manner compatible with the brain’s modular architecture.

## Acknowledgements

This work was supported by the Swiss National Science Foundation: Projects 31003A_182638 and 205320_215494/1, and Agreement 51NF40_180888. We thank Kristina Biedermann for setting up the human recordings.

