## Supplementary Materials for "Estimating the motor exploration in reinforcement learning"

### Animals

We used a total of 35 zebra finches (Taeniopygia guttata) that were bred and raised in our animal facilities in Zurich (Switzerland) and Orsay (France). The animals' ages at the beginning of the experiment in the range 90-200 days post hatch (dph). During the experiment, birds were individually housed in sound-attenuating recording chambers with a 14/10-hour day/night cycle. We used a total of $N = 20$ birds for LMAN lesions, from which $N = 8$ birds were pitch reinforced prior to lesioning LMAN. We reanalyzed pitch-reinforcement data in $N = 18$ birds from (Zai et al., 2024).All experimental procedures were performed in accordance with the Veterinary Office of the Canton of Zurich and with the French Ministry of Research and the ethical committee "Paris-Sud et Centre" (License number 2023-20).

### Song recording and pitch conditioning in birds

We recorded audio with a wall-attached microphone (Audio-Technica Pro4 and 2). The signal was amplified, filtered, and digitized at 32 kHz. We controlled sound acquisition and real-time pitch reinforcement with a custom LabView (National Instruments, Inc.) program (https://gitlab.switch.ch/hahnloser-songbird/cmatlab).

Pitch reinforcement: We trained a two-layer neural network to detect the beginning of a syllable containing a harmonic stack (Yamahachi et al., 2020). In detected syllables, we rigidly chose a suitable 16-24 ms segment and evaluated the fundamental frequency (pitch) as described in (Zai et al., 2020). When the pitch was either above or below a manually set threshold, we broadcast a 50-60 ms white-noise (WN) stimulus through a loudspeaker.

### Lesion surgery

We performed surgeries under general anesthesia (0.7-1.2% isoflurane, induction at 2%); providing analgesia with lidocaine delivered both globally (subcutaneous injection of 0.05 ml solution) and locally along the planned skin incision (0.05 g Emla® creme). We performed craniotomies above LMAN (1.7 mm lateral to the midline and 5.3 mm anterior to the confluence of sinuses, measured at a 35-degree angle of the flat anterior part of the skull). In each hemisphere, we assessed the center and extent of LMAN electrophysiologically using a single Tungsten wire electrode of 0.8-1.2 MΩ impedance (Micro Probe, Inc.).
We lesioned LMAN by pressure injecting (Picosprizter® III, Parker) ibotenic acid at 3-5 sites in LMAN (the center and 250 μm medially and laterally and/or anterior-posterior depending on electrophysiological response) via borosilicate glass pipettes (BF-120-69-10, Sutter instrument), as in (Zai et al., 2020).

### Histology

We euthanized 25 birds and perfused them with phosphate-buffered saline (PBS), followed by 4 % paraformaldehyde. We removed the cerebrum and kept the brain in paraformaldehyde for at least 24 hours before further processing. We embedded each hemisphere individually in agar before cutting it into 80-µm sagittal sections. We Nissl stained all slices with 0.3 % cresyl violet acetate solution.

### Quantification of lesion volume

We identified LMAN in sagittal brain slices by its large cell nuclei and its location in the nidopallium dorsal to Area X and between the lamina frontalis superior and the lamina pallio-subpallialis (Lovell et al., 2020; Nixdorf-Bergweiler & Bischof, 2007), see e.g. Supplementary Figure S1.

We estimated the lesioned LMAN volume as a fraction of the average intact LMAN volume in unmanipulated (control) adult males. To estimate the uni-hemispheric LMAN volume $V_{con}$ in control birds ($N = 8$), in each sagittal slice we quantified the area of LMAN and multiplied that with the section thickness. The summed volumes we averaged across ($N = 16$) hemispheres, yielding a reference LMAN volume of $V_{cont} = 0.12 \pm 0.02\ \mu m^3$, which is within the published range of $0.099\ \mu m^3$ (Scharff & Nottebohm, 1991) and $0.2\ \mu m^3$ (Nixdorf-Bergweiler et al., 1995). We quantified the unlesioned LMAN volume $V_{remaining}$ in each ($N = 17$) manipulated bird and hemisphere in the same way (see Supplementary Figure S1).

We then computed the fraction LMAN volume that was lesioned in the right hemisphere $F_{les,r}$ as the difference between the remaining LMAN volume and the reference volume, normalized by the reference volume:

$$F_{les,r} = \frac{V_{con} - V_{rem,r}}{V_{con}}.$$

We similarly computed the fraction $F_{les,l}$ of lesioned LMAN volume in the left hemisphere. In one hemisphere, the numerator was negative and so we set the fraction to zero.

The final fraction $F_{les}$ of lesioned LMAN volume in each bird we estimated as the average across the right and left hemispheres:

$$F_{les} = \frac{F_{les,r} + F_{les,l}}{2}.$$

The average fraction of lesioned LMAN volume was $\langle F_{les} \rangle = 61 \pm 28\%$ (std, $N = 17$ birds, range $0 - 100\%$, Supplementary Figure S2). We missed LMAN completely in both hemispheres in $N = 3$ birds. We included these 3 birds in the analysis because we used LMAN lesion volume as a predictor for the lesion effect. Excluding the $N = 3$ birds with 0% LMAN lesioned did not qualitatively change the results.

### Latent reinforcement learning equations

We assumed that the pitch $p_t$ of syllable rendition $t = 1, \dots, T$ is the sum of three sources of variability:

$$p_t = \epsilon_t + b_t + V_t, \tag{1}$$

where

- $\epsilon_t \sim P(\epsilon_t) = N(0, \sigma_\epsilon^2)$ are **explorations** attributed to the learner, forming independent and identically distributed (iid) Gaussian random variables with unknown variance $\sigma_\epsilon^2$,
- $b_t$ forms the learned **pitch bias**, and
- $V_t$ is the sum of **unreinforced motor primitives** (or 'other behavioral components').

The learner's **policy** $\pi_b$ is to generate an additive motor contribution $a = \epsilon + b$ with probability $\pi_b = p(a|b) = N(a - b, \sigma_\epsilon^2) = P(\epsilon)$, where we have omitted the time index $t$ for simplicity.

The learner performs policy gradient ascent (Williams, 1992). That is, it changes the policy $\pi_b$ along the gradient of a total **benefit function** $J$ composed of the expected reward $R$, offset by a cost proportional to the squared bias (to annihilate the bias is a simple model of song maintenance):

$$J(b) = \int da P(a|b) \left( R - \frac{\delta}{2\alpha} b^2 \right),$$

where $\delta$ and $\alpha$ are parameters ($\alpha$ we will interpret as the learning rate). As for the reward, we set $R_t = 0$ on escape trials and $R_t = -1$ on hit trials. We obtained similar results when we changed the definition of the reward to either $R_t = \{1,0\}$ or $\{1, -1\}$ on $\{escape, hit\}$.

The **gradient** $\nabla_b J$ is given by

$$\nabla_b J(b) = \int P(\epsilon) \epsilon R d\epsilon - \frac{\delta}{\alpha} b,$$

where we have used that $\epsilon = a - b$.

By updating the pitch bias in the direction of the policy gradient, $b \to b + \alpha \nabla_b J$, we obtain the following iterative update for the bias (after replacing the integral over the density $P(\epsilon)$ by samples):

$$b_t = b_{t-1} + \alpha R_{t-1} \cdot \epsilon_{t-1} - \delta b_{t-1}.$$

We add a small iid noise source $\eta_t$ with unknown variance $\sigma_\eta^2$ to this Equation (to perform parameter estimation using the formalism of a Kalman filter, see below), resulting in:

$$b_t = (1 - \delta) b_{t-1} + \alpha R_{t-1} \cdot \epsilon_{t-1} + \eta_t\ , \tag{2}$$

which is the **Pitch Learning Equation**.

We see that according to Equation (2), the learned motor bias $b_t$ performs a leaky correlation between explorations and rewards. The parameter $\delta$ acts as a time constant that defines how fast the bias decays towards zero (pitch maintenance). Without rewards ($R_t \equiv 0$), the bias acts like a noise source that is driven $\eta_t$; since we strictly impose $\delta < 1$, this noise source is non-white and so satisfies the presumed efficiency of reinforcement learning — only explorations are white).

The motor primitives $V_t$ in Equation (1) we kept modifying by trial and error until the **estimated explorations** $\hat{\epsilon}_t$ were close to ideal (i.e., the model was self-consistent, Figure 2e-i).

We obtained satisfactory self-consistency when using the motor primitives

$$V_t = p^* + c_t + d_t + o_t, \tag{3}$$

as detailed in the following.

**Song memory** $p^*$**:** During free unconstrained singing, syllable pitch has a stable mean $p^*$.

**Colored noise** $c_t$**:** We account for slow fluctuations in the pitch dynamics with a **colored noise component** $c_t$ that obeys a random walk: $c_t = (1-\tau)c_{t-1} + \varepsilon_t$, where $\tau$ is an unknown time constant and $\varepsilon_t \sim N(0, \sigma_\varepsilon^2)$ is a Gaussian random variable with unknown variance $\sigma_\varepsilon^2$. The autocorrelation function of colored noise decays exponentially with a decay constant set by the **time constant** $\tau$ and so for $\tau < 1$ is strictly different from the delta-function like autocorrelation of white noise.

**Circadian pattern** $d_t$**:** Zebra finch song exhibits circadian patterns manifest as repetitive pitch oscillations with a 24-h period. We modeled this pattern $d_t$ as a piecewise linear function constructed as follows.

We first introduced an auxiliary time function $h(t)$ equal to the time-of-day of trial $t$, expressed as a fraction of time since midnight ($0 \le h(t) < 1$). Next, we divided the daytime (during which the lights in the chamber are on) into $N_D = 5$ time periods $[H_i, H_{i+1}]$ ($i = 1, \dots, N_D$) such that on average (across all days) the same number of syllable renditions falls into each period (yielding more bins in the morning because birds tend to sing more in the morning). Thus, the time points $H_1, \dots, H_{N_D+1}$ (in fractions of a day) are such that the number of renditions in the interval $H_j \le h(t) < H_{j+1}$ are equal for all $j = 1, \dots, N_D$ (Fig. 2b). The daily pattern $d_t$ is then defined as

$$d_t = \sum_{j=1}^{N_D} D_j \theta_{h(t)}^j, \quad \text{where} \quad \theta_{h(t)}^j = \begin{cases} 1 - \frac{h(t)-H_j}{H_{j+1}-H_j} & \text{if } H_j \le h(t) < H_{j+1} \\ \frac{h(t)-H_{j-1}}{H_j - H_{j-1}} & H_{j-1} \le h(t) < H_j \\ 0 & \text{otherwise} \end{cases}$$

and where $D_j$ is a **circadian coefficient** corresponding to the circadian change in pitch at time $H_j$.

**History dependence** $o_t$**:** The pitch of a syllable is influenced by the history of singing, i.e. how much the bird recently sang. We assume that the number of times the bird produced a target syllable (i.e., a placeholder of the number of song motifs) in the last $x$ seconds has an additive effect on pitch. We modeled the history dependence of pitch as a fixed pitch offset

$o_t = O_{1+n_h(t)}$, where $n_h(t) \in \{1, \dots, N_h\}$ is the **number of target syllables within** $x$ **seconds preceding rendition** $t$.

We tested different history horizons $x$ (1 s, 2 s, 3 s, 5 s, 10 s, and 60 s) and different **history quantizations** $N_h$ used as a ceiling: $n_h(t) \to \min(n_h(t), N_h)$. We compared history model variants using a Bayesian information criterion (BIC) and found that the lowest average BIC was achieved for $x = 2$ seconds and $N_h = 5$ (average BIC = $2.5 * 10^5 \pm 1.1 * 10^5$, $N = 18$ birds, Fig. 2d), which we used throughout our experiments. For birds that produced at most 3 target syllables in the last 2 seconds, we set $N_h$ to $3 + 1 = 4$.

### The model written as a Kalman filter

Mathematically, latent RL models $p_t$ as a linear dynamical system with input $u_t$ that depends on some aspects of syllable renditions $t$ and $t-1$ and on the reward at $t-1$. To estimate the 16 model parameters $(p^*, \alpha, \delta, \tau, \sigma_\epsilon^2, \sigma_\varepsilon^2, D_t, O_k)$ and the hidden states $(\epsilon_t, b_t, c_t)$ of this model, we write it as a (linear) Kalman filter with time-dependent coefficients and inputs:

$$p_t = Hx_t + g_t \tag{4}$$

$$x_t = C_t \tilde{x}_{t-1} + q_t, \tag{5}$$

where $p_t$ (the pitch of rendition $t$) is the scalar obseration, $H = [1, 1]$ is the **observation matrix**, $x_t$ is the **latent state**, $C_t$ is the **transition matrix**, $\tilde{x}_{t-1} = \begin{bmatrix} x_{t-1} \\ u_t \end{bmatrix}$ is a combination of the latent state and a time-dependent **input vector** $u_t$, and $q_t$ and $g_t$ are **Gaussian i.i.d noises** with (diagonal) **covariance matrices** $Q$ and $G$. These noises need to be non-zero to estimate the parameters with the expectation maximization algorithm (see below). In practice, we fixed the output noise covariance $G$ to be very small, $g_t \sim N(0, 10^{-4})$. It follows that we estimated the explorations $\epsilon_t$ not as the output noise of the Kalman filter but as the hidden-state noise part of $q_t$.

Note, we also tested a version where the explorations $\epsilon_t$ correspond to the output noise $g_t$ and we obtained qualitatively similar results. However, this latter version requires some approximations (see Supplement). Note also it is possible to estimate parameters without the Kalman filter formulation (and without the addition small noise terms), but using nonlinear programming solvers.

**Baseline model**: To model baseline data (no reinforcement, $R_t \equiv 0$), we set the bias $b_t$ to zero. Equations (1-3) map to the Kalman filter Equations (4) and (5) with $x_t = \begin{bmatrix} c_t \\ p^* + d_t + o_t \end{bmatrix}$, $q_t = \begin{bmatrix} \varepsilon_t \\ \epsilon_t \end{bmatrix} \sim N(0, Q)$ with $Q = \mathrm{diag}(\sigma_\varepsilon^2, \sigma_\epsilon^2)$, and a time-dependent input vector $u_t = \left[ \left\{ \theta_{h(t)}^j \right\}_{j=1,\dots,N_D+1} \quad \left\{ \delta_t^j \right\}_{j=1,\dots,N_h} \right]'$ of dimension $N_D + 1 + N_h$. The transition matrix $C_t$ is not time dependent and given by $C = \sum_i \theta_i C^i$ with

$$(C^1)_{kl} = \begin{cases} 1 & \text{if } k = 1 \text{ and } l = 1 \\ 0 & \text{otherwise} \end{cases},$$

$$\left(C^{1+j}\right)_{kl} = \begin{cases} 1 & \text{if } k = 2, l = 1 + j \\ 0 & \text{otherwise} \end{cases}, \text{ for } j = 1, \dots, 1 + N_D + N_h,$$

and $\theta_i = \{(1-\tau), p^* + D_1, \ldots, p^* + D_{N_D+1}, O_1, \ldots, O_{N_h}\}$. We set $\theta_{2,\ldots,N_D+2}$ to $p^* + D_1, \ldots, p^* + D_{N_D+1}$ to enforce circadian patterns with zero average. On baseline data (before reinforcement starts or for LMAN-lesioned birds without reinforcement), we fitted model parameters $\theta_i$ and $(\sigma_\varepsilon^2, \sigma_\epsilon^2)$ as described in Section Parameter estimation.

**Reinforcement model**: We estimated the learning-related parameters $\Theta$ and $(\sigma_\epsilon^2, \sigma_\eta^2)$ by fixing the parameters that we estimated on baseline data except the exploration variance $\sigma_\epsilon^2$ that we estimated a second time for the purpose of testing the self-consistency of exploration estimation, Fig. 2n.

When we include the learner's bias $b_t$ (now nonzero) and fix the parameters estimated on the baseline data, the pitch values obey the Kalman filter equation with $x_t = \begin{bmatrix} b_t \\ c_t \\ p^* + d_t + o_t \end{bmatrix}$, $g_t \sim N(0, 10^{-4})$, $q_t = \begin{bmatrix} \eta_t \\ \varepsilon_t \\ \epsilon_t \end{bmatrix}$ with corresponding diagonal $Q$, and $u_t = \begin{bmatrix} c_t + d_t \\ c_{t-1} + d_{t-1} \end{bmatrix}$. The second dimension of $u_t$ is needed to estimate $\epsilon_{t-1} = (x_{t-1})_3 - (c_{t-1} + d_{t-1})$ appearing in the equation for $b_t$, where $(x_{t-1})_3$ stands for the third component of the vector $x_{t-1}$. Here, the transition matrix $C_t$ is time dependent because of the time dependence of reward ($R_t = \{-1,0\}$). The affected parameter is the learning rate $\alpha$. The transition matrix satisfies $C_t = C^0 + \sum_i \Theta_i C_t^i$ with $\Theta_i = \{(1-\delta), \alpha\}$ and

$$(C_t^1)_{kl} = (C^1)_{kl} = \begin{cases} 1 & \text{if } k = 1 \text{ and } l = 1 \\ 0 & \text{otherwise} \end{cases},$$

$$(C_t^2)_{kl} = \begin{cases} -R_{t-1} & \text{if } k = 1 \text{ and } l = 5 \\ R_{t-1} & \text{if } k = 1 \text{ and } l = 3, \\ 0 & \text{otherwise} \end{cases}$$

$$(C^0)_{kl} = \begin{cases} 1-\tau & \text{if } k = 2 \text{ and } l = 2 \\ 1 & \text{if } k = 3 \text{ and } l = 4. \\ 0 & \text{otherwise} \end{cases}$$

Note that there is no straightforward method to estimate all model parameters at the same time. When the learner's bias $b_t$ is nonzero, we would either have to add another iid noise term (i.e., one for exploration and the other for learning the parameters $D_t$ and $O_k$) or have to deal with nonlinearities arising from multiplicative parameter combinations (because $\alpha R_{t-1} \cdot \epsilon_{t-1}$ depends both on $\alpha$ and the parameters $D_t$ and $O_k$). Another possibility is to make simplifying approximations, i.e., to assume that the circadian rhythm does not change much between successive renditions (i.e., setting $D_t \approx D_{t-1}$) and that $\tau$ is small such that $(1-\tau) \approx 1$. When we tested fits using these simplifying approximations (see Supplement S2), we obtained qualitatively similar results compared to the results shown in the figures. Finally, another alternative would be to estimate parameters by setting $q_t^{(3)} \sim N(0, 10^{-4}), u_t = \begin{bmatrix} p^* + c_t + d_t \\ p_{t-1} \end{bmatrix}$, and $g_t \sim N(0, \sigma_\epsilon^2)$, in which case $\epsilon_{t-1}$ would be estimated as $p_{t-1} - r_t - c_t - d_t - o_t$. We also obtained similar results when trying this alternative, i.e., self-consistent fits with very low AMI.

### Parameter estimation

We estimate the parameters $\theta_i = \{(1-\tau), D_1, \dots, D_{N_d+1}, O_1, \dots, O_{N_h}\}$ and variances $(\sigma_\varepsilon^2, \sigma_\epsilon^2)$ governing the baseline period and the parameters $\Theta_i = \{(1-\delta), \alpha\}$ and $(\sigma_\eta^2, \sigma_\epsilon^2)$ of the latent learner from behavioral data using the iterative expectation maximization (EM) algorithm. The EM objective is to maximize the log-likelihood of the data given the model parameters (Ghahramani & Hinton, 1996; Shumway & Stoffer, 1982).

The standard EM algorithm works well when the time-dependent input $u_t$ depends on explicit variables (Cheng & Sabes, 2006) such as the time of renditions $t$ and $t-1$. However, in our case there is also a time-dependent transition matrix $C_t$ with components that are either zero (e.g. $(C^1)_{kl} = 0$ except if $k = l = 1$) or are shared among model components (e.g. $(C_t^2)_{13} = -(C_t^2)_{15}$ in the reinforcement case). To deal with this situation, we used the method described in (Cheng & Sabes, 2006; Ghahramani & Hinton, 1996; Shumway & Stoffer, 1982) that applies when the following holds:

1. $Q$ is diagonal
2. $C_t$ can be written as $C_t = C_t^0 + \sum_i \Theta_i C_t^i$ with $\left(C_t^i\right)_{m,n} = 0 \text{ for } m \neq m_i$, where $m_i$ is the row of $C_t$ in which parameter $\Theta_i$ is located

For more details and the EM algorithm, see Supplement.

Before estimating the parameters, we perform mean subtraction (we subtract $\bar{p} = \frac{1}{T_b}\sum_{t=1}^{T_b} p_t$ from the measured pitch values, where $T_b$ marks the end of baseline. This is not crucial but helps to avoid numerical instabilities arising from the large ratio of $10^6$ between the smallest parameter (e.g. $\delta \approx 0.001$) and the largest parameter ($p^* \approx 1000$ Hz). Thus, in practice, the estimated target pitch is $\hat{p}^* = \bar{p} + \frac{1}{T_b}\sum_{t=1}^{T_b} \hat{d}_t$, which we keep constant during reinforcement.

We further add the constraint that the history dependence averages to zero and does not add an overall pitch bias; we enforce this constraint after every EM iteration by transferring the non-zero history average to the circadian pattern: $D_i \to D_i + \frac{1}{T_b}\sum_{t=1}^{T_b} o_t$ and $O_i \to O_i - \frac{1}{T_b}\sum_{t=1}^{T} o_t$.

### Fit of baseline vs WN exploration variance

We fitted the BL vs RL exploration variances in Fig. 3f using Matlab's regress function, determining the p value at which the unit slope $m = 1.0$ fell outsize the estimated confidence interval, which amounts to a Student's t-test.

### Mutual average information

Unless specified otherwise, we calculated the mutual average information (MAI) on 50 log-spaced rendition lags (integers) from $10^0$ to $10^3$ using the MATLAB (Mathworks Inc) function mai (https://www.mathworks.com/matlabcentral/fileexchange/880-mutual-average-information). To account for the non-zero MAI of a finite time series, we computed the average MAI of three randomly drawn Gaussian iid. time series of the same length as the pitch trajectory in each bird (plotted as dashed lines in Fig. 2h). We then normalized the MAI for all birds individually such that zero corresponds to the average MAI of the three random time series and one to the bird's MAI at lag 1 (Fig. 2i).

### The Kalman filter used as a generative model

We validated the fitted Kalman model in terms of the artificial data it generates. For each bird and corresponding fitted parameters, we let the model generate the same number of pitch renditions as produced by the bird, using the renditions' time stamps. As we did in the experiments, for each simulated day, we set the reward threshold to the median of the simulated pitch values of the previous day. We simulated each birds' trajectory 100 times; these we compared with the true trajectory after smoothing both simulations and true trajectories via a running average of the last 50 renditions (see Fig. 2j for an example).

For each smoothed simulation $i$ ($i = 1, \dots, 100$), we calculated the average **pitch standard deviation** $E_i$, i.e., the root-mean square (RMS) pitch deviation from the mean simulated pitch trajectory. We also computed the RMS pitch deviation $E_0$ of the observed pitch trajectory from the simulated mean, i.e., the bias of the mean simulated trajectory (black bar in Fig. 2k).

To compute the **normalized RMS pitch error** $E_n$ (Fig. 2l), we averaged the stimulated RMS deviations $E = \langle E_i \rangle_i$ and divided $E_0$, by this average, $E_n = E_0/E$. The value $E_n = 1.0$ (see Results) means that the true trajectory is as far away from the stimulated mean as is the typical simulated trajectory, i.e. that the observed trajectory is not an outlier.

To compare the latent learner model with traditional RL models where all behavioral variability is exploited during learning, we simulated a classical RL model in which we replaced $R_{t-1} \cdot \epsilon_{t-1}$ in Equation (2) by $R_{t-1} \cdot (\epsilon_{t-1} + V_{t-1}) = R_{t-1} \cdot (\epsilon_t + c_t + s_t + o_t)$, Fig. 2l (right). We compared the latent with the classical RL model variants in terms of their distributions of normalized RMSEs via a paired two-sided t-test (we tested whether their average normalized errors are equal). The classical RL model produced normalized RMSEs across birds averaging to $E_n = 1.8$, which was rejected as a model of the observed data ($p = 2 * 10^{-4}$, t-test for $H0$: $E_n = 1$, $df = 17$): A value $E_n = 1.8$ means that the true trajectory is separated from the average simulated trajectory by more than 92% of simulated trajectories. i.e. the observed trajectory lies in the tail of the simulated distribution and is unplausible.

### Estimation of exploration variance

To test how much data is needed to estimate the exploration variance and whether it is possible to do so only during baseline (no RL), we estimated the baseline model parameters (including $\sigma_\epsilon^2$) using either 1, 3, or 5 days of baseline data (7 birds were excluded for 5-day estimation because of data unavailability). We then compared the estimated exploration variance during baseline $\hat{\sigma}_{BL,\epsilon}^2$ to the estimated variance $\hat{\sigma}_\epsilon^2$ during and after RL (excluding baseline, as in Supplement S2).

Across birds, the Pearson correlations $\rho$ between $\hat{\sigma}_{\mathrm{BL},\epsilon}^2$ and $\hat{\sigma}_\epsilon^2$ were large regardless of the number of baseline days included (Fig. 2n):

- 1 day: $\rho = 0.89$, $p = 8.2 * 10^{-7}$ ($N = 18$ birds)
- 3 days: $\rho = 0.91$, $p = 1 * 10^{-7}$ ($N = 18$ birds)

- 5-days: $\rho = 0.91$, $p = 9.9 * 10^{-5}$ ($N = 11$ birds)

### Human subjects

We recruited $N = 18$ human subjects (13 females, 5 males) at the University of Zurich, aged between 18 and 60 years. All experiments procedures were performed in accordance with the Research Ethics Committee of ETH Zurich (2017-N-52). $N = 4$ subjects were excluded (3 females and 1 male) because of unstable pitch (bimodal distribution). None of the participants reported to be either a professional singer or having absolute pitch. $N = 3$ participants were recorded before the other participants in a preliminary study using a slightly different experimental design (indicated below). We did not see any systematic difference in those participants and thus decided to include them in the analysis (total $N = 14$ subjects); when excluding those subjects we obtained qualitatively similar results.

### Pitch conditioning in humans

Audio signals were acquired with a microphone (SM6 model from Røde) placed in front of the subjects. The signals were amplified, filtered and digitized at 32 kHz with the same system we used for birds. Subjects were instructed to repeat the syllable "Da" at a speed of about 2 "Da's" per second. For all except the first 3 participants, a metronome was shown on the screen to help them maintain a steady pace. We detected the onset of a syllable by thresholding sound amplitude (RMS sound waveform), the threshold was kept constant at a level well above the noise level of the room.

During a test session, subjects were instructed to adjust the sound amplitude of their voice (displayed on the screen) such that it fell below the threshold between vocalizations and by a factor of 4-6 above the threshold during the vocalizations. We evaluated pitch in a 16-ms window at a fixed latency of 88 ms to vocalization onsets.

The experiment was conducted in short sessions interleaved by breaks. During the breaks, participants were encouraged to drink water and/or take a lozenge for the throat.

Participants performed first 1-3 baseline sessions of roughly 6 minutes each and then 1-2 conditioning sessions (the first 3 participants performed baseline and conditioning sessions on different days). Before each conditioning session, participants were asked to roll a dice to determine the initial white-noise contingency (1-3: WN on low pitch; 4-6: WN on high-pitch). On the second conditioning session we inverted the contingency independent of their second roll.

During a conditioning session, we first recorded 1 minute of data without conditioning, then 3 minutes with white noise conditioning, and again 2 minutes without conditioning (in one participant we omitted a white-noise free period after conditioning). We set the white-noise threshold to the median pitch of the 20 most recent renditions. We automatically adjusted the threshold when the escape rate during the last 20 renditions was above 80%. We also adapted the threshold when the escape rate was below 5%, to prevent subjects from getting stuck at pitch values where WN becomes uninformative (where all renditions trigger WN).

Participants were given the following initial instructions: "During the third and fourth sessions, you will sometimes hear a white noise sound after your Da. When you hear the white noise sound, please try to stay as calm as possible and continue saying Da as before. The goal is to avoid this white noise sound but stay as relaxed as possible." The experimenter verbally repeated the instructions before the second conditioning session.

Humans shifted their pitch away from the WN zone as expected: the pitch difference between the first and last 50 utterance renditions of a session was $d' = 4.8 \pm 7.8$ ($p = 0.02$, one-tailed t-test, $tstat = 2.30$, $df = 13$), and similarly the pitch of the last $50$ renditions differed from the pitch of the corresponding $50$ renditions of the preceding baseline session by $d' = 5.1 \pm 7.9$ ($p = 0.03$, two-tailed paired t-test, $tstat = 2.43$, $df = 13$, see Methods, Fig. 4c). After cessation of WN delivery, during the short recovery time period we provided, subjects tended to revert their pitch towards baseline by $d' = -1.5 \pm 3.3$ ($p = 0.07$, one-sided t-test, $tstat = -1.62$, $df = 12$, excluding one subject who was not given a recovery time period).

To estimate explorations, we simulated a model without circadian pattern and history dependence because of the brevity of the baseline and WN sessions. Because of considerable pitch variability among sessions, we analyzed explorations merely in the last baseline session. For all subjects except two, we analyzed the first conditioning session because during the second conditioning session subjects tended to get distracted or start to play around with the system by strongly modulating their voice. Two subjects did not understand the task correctly in the first conditioning session, which is why we analyzed data from their second session.

To quantify pitch changes in either the baseline, reinforcement, or recovery period (Figure 4c), we compared the average pitch of the last 50 renditions to the average pitch of the first 50 renditions in that period. The reinforcement period corresponds to the trials in the conditioning session on which the threshold was different from zero and the recovery period corresponds to all renditions on which the threshold was set back to zero (participants could not trigger white noise any more).

We also compared pitch changes during baseline and conditioning periods at identical rendition lags since beginning of the period (i.e. if the first 50 renditions during reinforcement were renditions 201-250, then the same rendition lags were used to average baseline pitch). We tested for differences in average pitch across periods using paired one-tailed t-tests over subjects and we tested for nonzero pitch changes during reinforcement using two-tailed t-tests. We tested for correlations between exploration variances associated with baseline and WN sessions in terms of the Pearson correlation coefficient.

### Pitch steps

We detected discontinuities or steps in human pitch dynamics (Supplementary Figure S3) when the average of the $n_s = 20$ pitch values before a given rendition differed from the average of the $20$ pitch values following (and including) that rendition by more than a factor $\Theta$ of the average pitch across all 41 renditions: A step at rendition $i$ was detected when the pitch step satisfied $|p_i^l - p_i^r| > \Theta p_i^m$, where $p_i^l = \langle p_i \rangle_{[i-20,i]}$, $p_i^r = \langle p_i \rangle_{[i,i+20]}$, $p_i^m = \langle p_i \rangle_{[i-20,i+20]}$. We detected discontinuities using $\Theta = 0.05, 0.01$, or $0.1$.

Once we detected a discontinuity, we removed it from the pitch trajectory to look for more discontinuities until none was left in the given session. We iteratively eliminated the largest pitch discontinuity in terms of the maximum pitch step. Elimination was done by adding the step size to all subsequent renditions: if the largest discontinuity was at rendition $i$, then we applied the transformation $p_{j>i} = p_{j>i} + |p_i^l - p_i^r|$ to all following pitches, see Supplementary Figure S3.

We then analyzed the rate of pitch discontinuities, in particular the rate $rA$ following WN experience: A pitch discontinuity followed WN experience if WN was delivered on any of the 20 renditions preceding the discontinuity. We calculated this discontinuity rate as $rA = \frac{nD_{WN}}{T}$, where $nD_{WN}$ is the number of discontinuities following WN experience and $T$ the number of pitch renditions in the session (we deliberately ignored discontinuities at session boundaries).

We compared $rA$ to the chance-level rate $rC$ that assumes no relationship between WN and discontinuities. This latter rate we calculated as $rC = \frac{nWN}{T}$, where $nWN$ is the number of renditions preceded by WN. We compared the rates $rA$ and $rC$ using a test for equality of proportions (i.e., the rates are probabilities per rendition) using the $Z$ test statistics defined as $Z = \frac{r_A - r_C}{\sqrt{r(1-r)(\frac{1}{T}+\frac{1}{n_D})}}$, with $r = \frac{nD_{WN}+n_{WN}}{nD+T}$ and with $nD$ the number of discontinuities in that subject. A subject responded to WN by significantly when the difference between their baseline and WN-associated discontinuity rates satisfied $|Z| > 1.96$, corresponding to $p < 0.05$. Subjects that responded to WN by increasing the pitch step rate were said to explore stepwise.

We report results for $n_s = 20$ and $\Theta = 5\%$; the results were quite robust to changes of these parameters, detailed in the following.

- For $\Theta = 10\%$ ($n_s = 20$) we observed fewer pitch steps: 3 subjects showed a stepping rate that differed between WN and baseline (plus one subject showed a mere trend: $Z = 1.67$), all four subjects increased their stepping rate during WN, all in agreement with directed stepwise exploration.
- For $\Theta = 2.5\%$ ($n_s = 20$) four subjects showed a stepping rate that different between WN and baseline (plus one subject showed a trend: $Z = 1.45$): Four subjects increased the stepping rate during WN, and one decreased the stepping rate ($Z = -2.9$). Note that at such a small $\Theta$, it can be argued whether the detected steps are true discontinuities, since they are large numbers of them, without showing a clear change in pitch level.
- For $n_s = 10$ ($\Theta = 5\%$), 5/14 subjects showed a significantly different stepping rate during WN, all of them increased the stepping rate, in agreement with directed stepwise exploration.
- For $n_s = 10$ ($\Theta = 5\%$), 6/14 subjects significantly changed the pitch step rate in response to WN ($|Z| > 1.96$), five produced more pitch steps (all in the direction of WN escape), one decreased the pitch step rate ($z = -2.1$). The number of steps was in general too small to assess the significance of whether they were aligned with the direction of WN escape or not, we did not observe any clear trend either.

None of the birds produced pitch steps at the level of $\Theta = 5\%$. At the finer level of $\Theta = 2\%$, we detected some pitch steps. Namely, three of 18 birds significantly changed the pitch stepping rate after experiencing WN ($|Z| > 1.96$), two of them with a higher stepping rate during WN, and one with a lower stepping rate, providing little evidence overall of WN-directed stepwise exploration in Birds. Nevertheless, in three birds, the WN-associated steps were biased towards WN escape, suggesting that birds are capable of stepwise exploitation, in line with previous reports (Costalunga et al., 2023; Veit et al., 2021; Zai et al., 2024). Thus, human pitch learning might make use of a similar learning mechanism driven by ideal explorations.
